# The persistence of Nontypeable *Haemophilus influenzae* fuels airway inflammation

**DOI:** 10.1101/2020.05.25.107995

**Authors:** Fabio Saliu, Giulia Rizzo, Alessandra Bragonzi, Lisa Cariani, Daniela M. Cirillo, Carla Colombo, Valeria Daccò, Daniela Girelli, Sara Rizzetto, Barbara Sipione, Cristina Cigana, Nicola I. Lorè

## Abstract

Nontypeable *Haemophilus influenzae* (NTHi) is commonly isolated from airway of cystic fibrosis (CF) patients. However, to what extent NTHi persistence contributes to the lung inflammatory burden during CF chronic airway disease is controversial. Here, we aimed at determining the pathological role of NTHi persistence in a cohort of CF patients and in a newly generated mouse model of NTHi persistence.

In our study cohort, we found that CF patients chronically colonized by NTHi had significantly higher levels of IL-8 and CXCL1 than those who were not colonized. To better define the impact of NTHi persistence in fuelling inflammatory response, we developed a new mouse model using both laboratory and CF clinical strains. NTHi persistence was associated with chronic inflammation of the lung, characterized by recruitment of neutrophils and cytokine release (KC, G-CFS, IL-6 and IL-17A) at 2 and 14 days postinfection. An increased burden of T cell mediated response (CD4^+^ and γδ cells) and higher levels of matrix metalloproteinase 9, known to be associated with tissue remodelling, were observed at 14 days post-infection. Of note we found that both CD4^+^IL-17^+^ cells and levels of IL-17 cytokine were enriched in mice at advanced stage of NTHi chronic infection. Moreover, by immunohistochemistry we found CD3^+^, B220^+^ and CXCL-13^+^ cells localized in bronchus-associated lymphoid tissue-like structures at day 14.

Our results demonstrate that NTHi persistence exerts a pro-inflammatory activity in the human and murine lung, and could therefore contribute to the exaggerated burden of lung inflammation in CF patients.

## • Introduction

Recurrent or chronic infections by several opportunistic pathogens are common in patients affected by different chronic respiratory diseases, including chronic obstructive pulmonary disease (COPD) and cystic fibrosis (CF) [1–3]. Infections by *Nontypeable Haemophilus influenzae* (NTHi), a non-capsulated Gram-negative bacterium, are associated with worse clinical prognosis in several chronic respiratory diseases, such as COPD [4, 5], while its pathogenic potential in CF disease is still controversial.

CF is an autosomal recessive genetic disease caused by mutations in the CF transmembrane conductance regulator (CFTR) gene [4]. A deficiency of the CFTR protein leads to an alteration of the ionic transport, production of thick secretions and obstruction of the gland ducts with progressive epithelial damage. CF patients display an impaired mucociliary clearance in the airways that favours bacterial colonization and/or persistent infections early in childhood. Vicious cycles of chronic inflammation and infections have been associated with immunopathology and decline in pulmonary function [6, 7]. The natural history of airway colonization in children with CF is initially characterized by the isolation of NTHi, *Streptococcus pneumoniae* and/or *Staphylococcus aureus* infection, followed by other pathogens such as *Pseudomonas aeruginosa* [8–10]. The pathogenic potential and contribution of *P. aeruginosa* or *S. aureus* to the CF morbidity has been deeply described and demonstrated in details [11, 12]. As to NTHi pathogenicity in CF, it has been reported that the presence of this pathogen can be associated with acute exacerbation episodes [13]. However, there are no observations on the effect of NTHi persistence in the progression of CF chronic airway diseases, due to the presence of several clinical and microbiological cofounding factors, such as concomitant infections [8, 14] and lack of available human cohorts. Previous studies on NTHi were mainly focused on the acute virulence of genotypically and phenotypically different isolates [15–17] rather than on the pathogenic potential of NTHi persistence in the airways. The unavailability of appropriate animal models represents another limitation for the studies on the potential contribution of NTHi persistence to CF lung disease.

To validate our hypothesis that NTHi persistence contributes to fuelling the CF lung inflammatory burden, we have measured the concentrations of a few pro-inflammatory cytokines in respiratory samples from CF patients colonized by NTHi alone or not colonized by any other CF opportunistic pathogens. In addition, NTHi laboratory and clinical strains were used to test their pathogenic potential i) in a new mouse model for NTHi long-term infection mirroring clinical lung persistence and ii) in *in vitro* host-pathogen experiments. Overall, our data demonstrate that NTHi persistence, independently from NTHi phenotypic diversity, may contribute to fuelling the exaggerated burden of lung inflammation in CF patients.

## • Methods

### Ethical committee

Animal studies were performed according to the protocols set forth by the Italian Ministry of Health guidelines for the use and care of experimental animals (IACUC protocols #920) and approved by the San Raffaele Scientific Institute (Milan, Italy), Institutional Animal Care and Use Committee (IACUC). Human samples and the bacteria from CF patients were collected at the Cystic Fibrosis Center of Milan, Italy. The study was approved by the Ethical Committees of San Raffaele Scientific Institute and of Fondazione IRCCS Ca’ Granda, Ospedale Maggiore Policlinico, Milan, Italy. Written informed consent was obtained from patients enrolled, or their parents, according to the Ethical Committee rules, in accordance with the laws of the Italian Ministero della Salute (approvals #1874/12 and 1084/14).

### Human respiratory samples and lung function

Nasopharyngeal aspirates from 19 CF patients with variable genotypes were collected from the Regional CF Center at Milan’s Ospedale Maggiore Policlinico during routine care visits as described in supplementary methods 1. Seven patients (five females) with an age range between 13-20 years were negative for CF opportunistic pathogens, 12 patients (five females), aged between 4-9 years, did not have opportunistic pathogens, but NTHi colonization.

### Bacterial Strains

ATCC 49766 is a reference laboratory strain purchased from ATCC. The eight NTHi clinical strains tested in *in vivo* or *in vitro* were provided by CF Microbiology Laboratory, IRCCS Ca’ Granda, Ospedale Maggiore Policlinico (Supplementary Table S1).

### Mouse model of chronic NTHi lung infection

Persistent airway infection was obtained by intratracheal injection of agar beads containing ~1 × 10^7^ colony forming units (CFU) of NTHi (ATCC 49766 or NTHi 50, a CF clinical strain) or empty beads (as sham control) in 20-22g C57BL6/NCrl male mice (8-10 weeks, Charles River). Agar beads were produced by refining a previously described protocol [12, 18, 19] as described in Supplementary methods. Animals were euthanized after 2, 7 and 14 days by CO2 administration and lungs were recovered. Sample processing was performed as described in Supplementary methods.

### Flow cytometry and intracellular cytokines staining

After euthanasia, lungs were harvested and mechanically dissociated in GentleMACS C tubes (Miltenyi Biotec) using gentleMACS dissociators, followed by straining through a 70-μm filter. Antibody staining was performed as previously described [19, 20]. Briefly, lung cell suspensions (1.5 × 10^6^ cells) were incubated with blocking buffer (5% rat serum and 95% culture supernatant of 24G2 anti-FcR mAb-producing hybridoma cells) for 10 min at 4 °C. Then, cells were stained for 20 min at 4 °C in the darkness with different combinations of antibodies (listed in Supplementary tables 2, 3 and 4). For intracellular cytokine staining, 1–3 × 10^6^ cells were stimulated with PMA (3 ng/mL) and Ionomycin (1 μg/mL) for 4 h at 37 °C (the last 2 h with Brefeldin A (5 μg/mL). Cells were surface stained, fixed (paraformaldehyde 2% for 10 min), permeabilized (2% FBS, 0.2% NaN_3_, 2% rat serum, 0.5% saponin in DPBS) and then stained for intracellular cytokines for 30 min at RT in darkness. Acquisition and analyses were performed using fluorescence-activated cell sorting FACSCanto cytometer (BD Biosciences) and FlowJo Software (Tree Star), as previously described [21].

### Histological analysis

Lungs were removed and fixed in 4% paraformaldehyde. After paraffin embedding, consecutive 5 μm sections from the middle of the five lung lobes were used for histological and immunohistochemical examination. Sections were stained with Hematoxylin and Eosin (H&E) or immunostained as previously described with anti-CD3, anti-B220, anti CXCL-13, anti F4/80 antibodies [22]. A description of the primary antibodies, together with dilutions and unmasking techniques, is provided in the Supplementary table 5. Images were acquired using the Leica Biosystems Aperio CS2 scanner and analysed with the Aperio ImageScope software.

### Morphometric analysis

A quantitative analysis was performed on tissue slides by using the Aperio ImageScope software. Lymphoid aggregates were defined as cell clusters positive for the CD3 molecule and were quantified as number of clusters per cm^2^ of lung.

### ELISA

Expression of cytokines/chemokines in lung homogenates was assessed by ELISA using DuoSet kits (R&D Systems, Minneapolis, MN) for growth-regulated alpha protein (CXCL1/KC), C-X-C motif chemokine 2 (CXCL2/MIP-2), granulocyte colony-stimulating factor (G-CSF), interleukin-6 (IL-6), interleukin-17A (IL-17A) and pro-matrix metalloproteinase 9 (Pro-MMP9) accordingly to manufacturer’s instructions.

### NTHi infection in A549 cell line

Human alveolar epithelial A549 cells were cultured and challenged with NTHi strains as described in the supplementary Methods. Cell viability was assessed by measuring the lactate dehydrogenase released in the supernatant with CytoTox 96 Non-Radioactive Cytotoxicity Assay (Promega) following manufacturer’s instruction.

### Statistical Analysis

Statistics were performed with GraphPad Prism for statistical computing. Data at a specific time-point were compared through a nonparametric two-tailed Mann-Whitney test. Incidences of mortality and chronic colonization were compared using Fisher exact test. Statistical analyses were considered significant at p < 0.05.

## • Results

### Impact of NTHi colonization on human CF lung inflammation

We evaluated the levels of CF pro-inflammatory cytokines (IL-8 and CXCL1) in respiratory samples of 19 CF patients with NTHi colonization or uncolonized by NTHi. (See material and Methods). We found that CF patients colonized by NTHi had significantly higher levels of both IL-8 and CXCL1 than those uninfected (Figure 1 a, b). These data suggest that NTHi persistence may contribute to the CF inflammatory burden.

**FIGURE 1.**
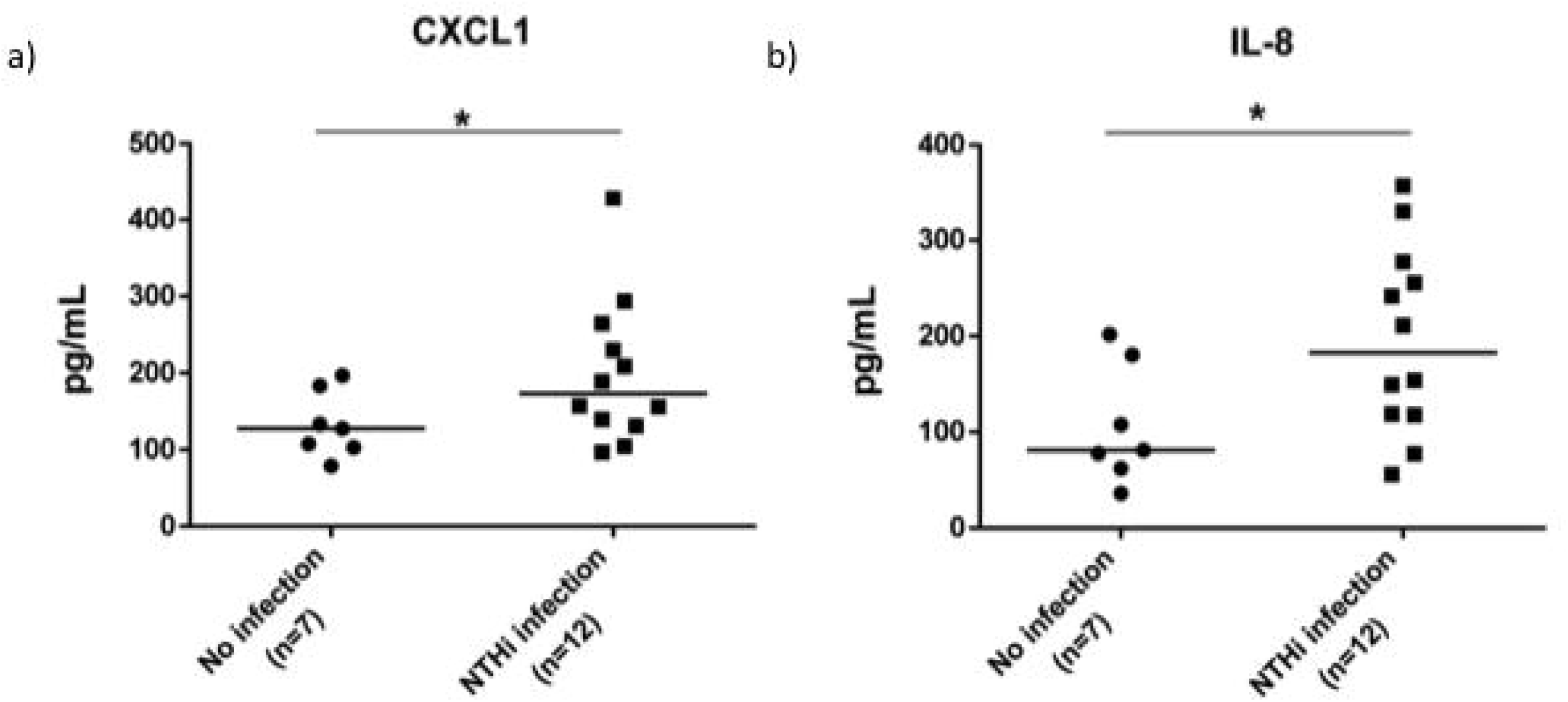
Contribution of NTHi persistence to lung inflammatory burden in cystic fibrosis patients. Levels of proinflammatory chemokines (a) CXCL1 and (b) IL-8 were evaluated by ELISA in respiratory samples from a retrospective study in CF patients colonized exclusively by NTHi (n=12) or uncolonized (n=7) by any recognized CF pathogen. *p<0.05 compared to control (Mann-Whitney t test).

### Development of a new murine model to mirror NTHi persistence as in human pathology

To investigate whether NTHi colonization can induce an inflammatory response, we developed a new murine model that mirrors NTHi persistence as observed in human CF pathology.

We adapted a murine model, previously developed for chronic bacterial lung infections [19] [12], to study NTHi persistence over the long term in lower-middle airways. We obtained a NTHi colonization with the 49766 reference strain up to 14 days with bacterial loads of ≈ 10^6^ CFU/lung (Figure 2 a) and with a high incidence of chronic infection (80 %) at 14 days post-infection (Figure 2 b). It is worth noting that NTHi was confined to the lung compartment and did not spread systemically, as inferred by absence of colonization in the spleens of infected mice (data not shown).

**FIGURE 2.**
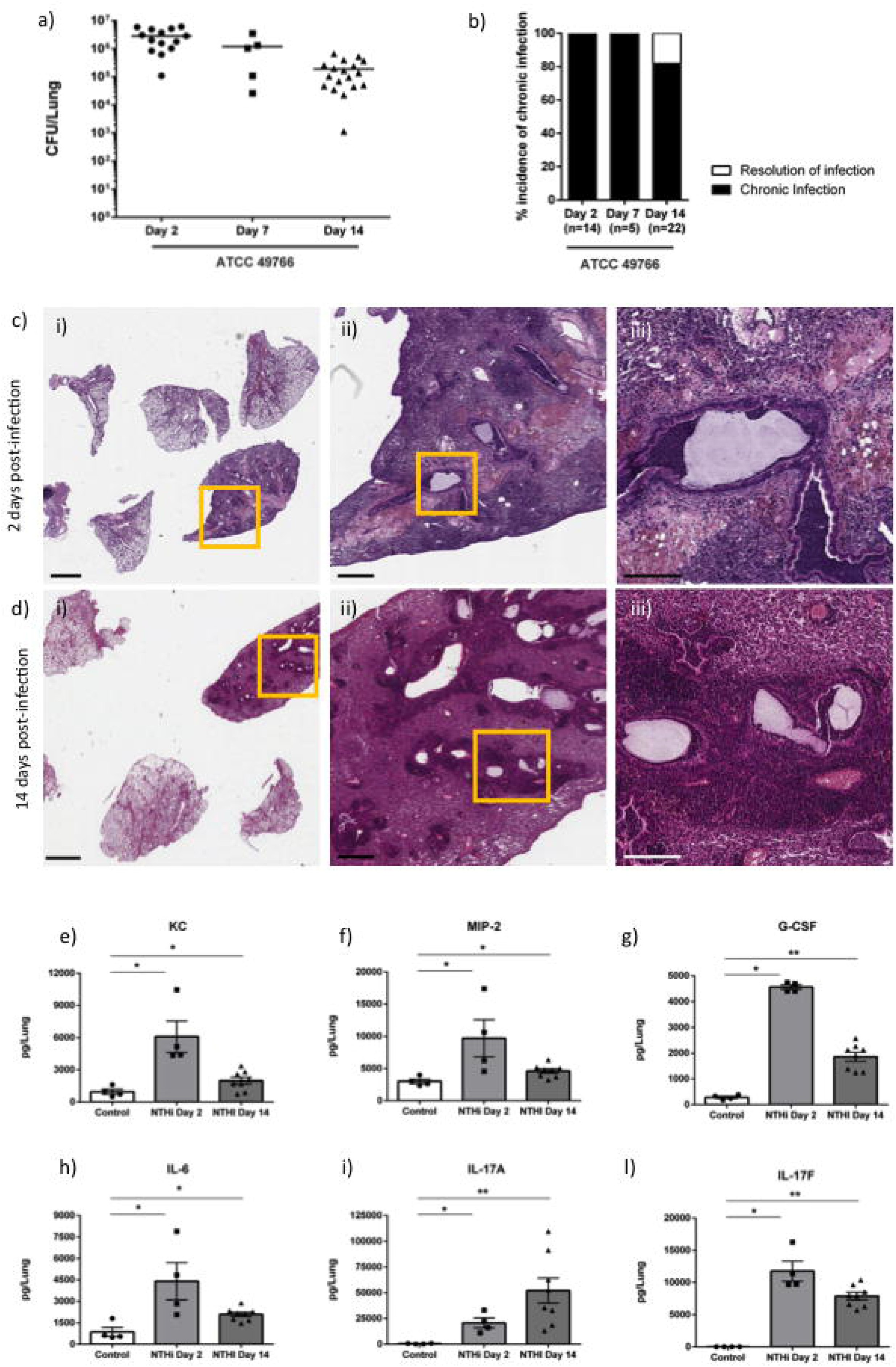
Bacterial burdens, histopathology and pro-inflammatory cytokines during the NTHi persistence in murine airways. C57BL/6N mice were infected intratracheally with NTHi embedded agar beads. Bacterial loads in the lung (a) and the incidence of chronic infection (b) were measured at 2, 7 and 14 days post infection. H&E staining of lung tissue sections was performed at 2 (c) and 14 day d) post infection. Representative images are shown from the lower i) to the higher iii) magnification. i) scale bar=2 mm; ii) scale bar=500 μm; iii) scale bar=200 μm. KC (e), Mip-2 (f), G-CSF (g), IL-6 (h) and IL-17A (i) expression in the lung of the infected mice was assessed by ELISA at both 2 and 14 days post infection. The data are pooled from at least two independent experiments (n=2-7). *p<0.05, **p<0.01, ***p<0.001 compared to control (Mann-Whitney t test).

Focusing on the early (2 days) and advanced (14 days) stages of infection, we evaluated histopathology and the levels of pro-inflammatory cytokines related to neutrophilic recruitment into the lungs. H&E staining on day 2 and 14 days post-infection localized the agar beads to bronchial lumens. In term of inflammation, the early phase was mainly characterized by the acute neutrophil and macrophage recruitment (Figure 2 c), while the long-term NTHi persistence was featured also by adaptive immune response including the formation of organized macro structure of lymphocytes (Figure 2 d) similar to those observed for chronic *P. aeruginosa* infection [23] [19]_[12].

The levels of KC, MIP-2, G-CSF, IL-6, IL-17A and IL-17F were significantly increased in comparison to control mice at 2 or 14 days post-infection, reflecting the higher recruitment of innate immune cells described in the histopathological analysis (Figure 1 e,f,g,h,i). The production of KC, MIP-2, G-CSF, IL-6 and IL-17F were reduced at day 14, despite remaining significantly higher than those found in sham infected (control mice). Differently, IL-17A levels increased by day 2 and remained high over the course of NTHi infection, suggesting the involvement of type 17 immunity. Overall, this murine model for NTHi persistence can mirror the acute/early and advanced stages of chronic lung infection observed during the course of human airway diseases.

### NTHi persistence in the lung promotes sustained type 17 immunity and remodeling processes

In order to dissect the lung infiltrating cells, FACS analysis was performed. The number of infiltrating leukocytes was significantly higher in samples recovered from infected mice at both day 2 and 14 post-infection when compared to control mice (Figure 3 a). Neutrophils and monocytes were recruited in infected lungs at both time points (Figure 3 b, c), while alveolar macrophages and dendritic cells (DC) were increased only at 14 days post infection (Figure 3 d, e). On the other hand, neutrophil and monocyte counts decreased on day 14 post-infection, although they remained higher than in control mice. Additionally, T cells were significantly more represented among infected mice compared to control ones (Figure 3 f). While both ^+^ and CD4^+^ T cell subsets were significantly enriched both at day 2 and day 14 postinfection (Figure 3 g, h), CD8^+^ T cell subsets only increased at day 14 (Figure 3 i).

**FIGURE 3.**
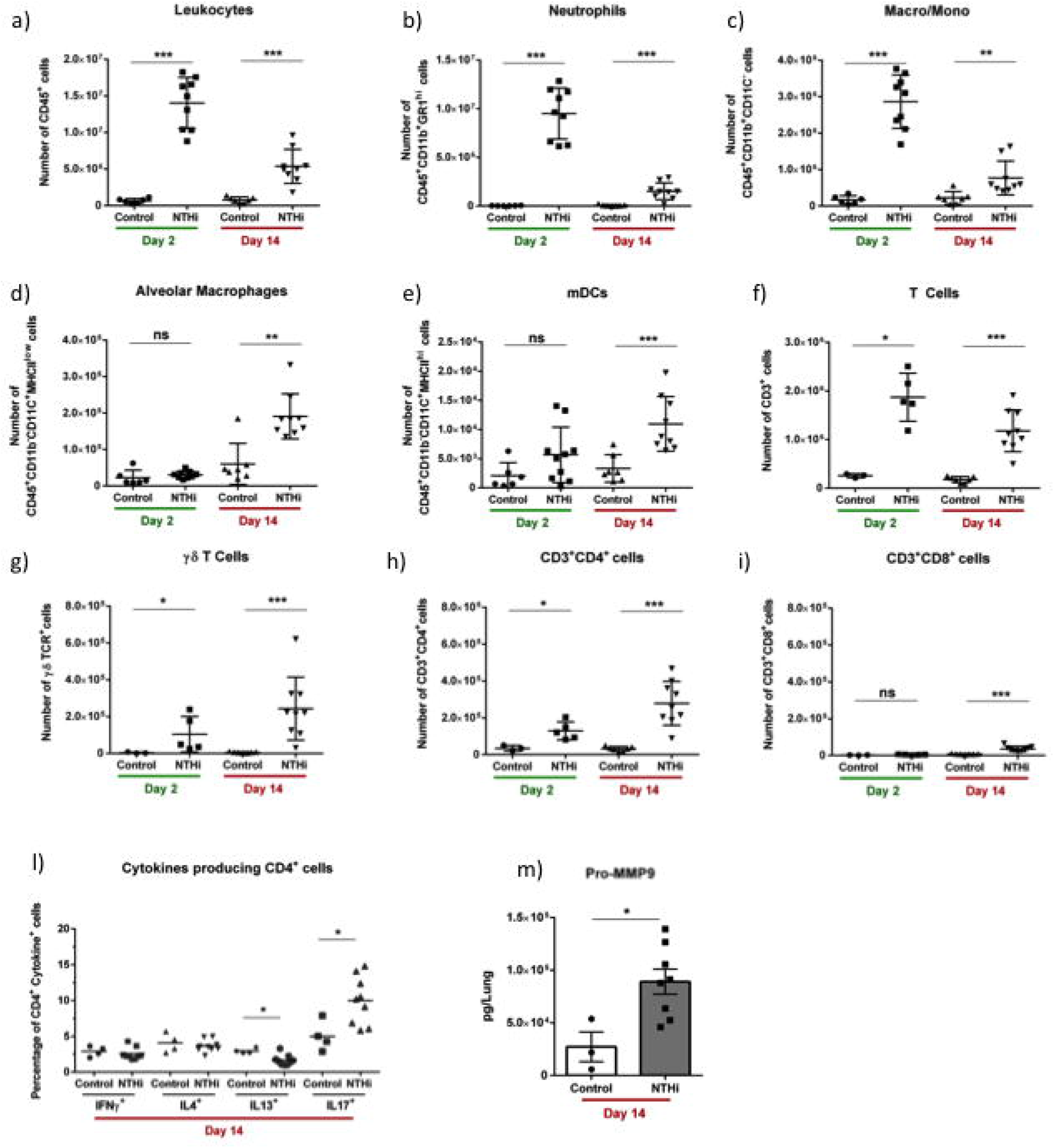
Immune response during the development of NTHi persistence in mice. Mice were infected intratracheally with NTHi embedded agar beads and lungs were dissociated at 2 and 14 days post infection. Flow cytometry was performed to quantify the total number of (a) leukocytes, (b) neutrophils, (c) monocytes/macrophages, (d) alveolar macrophages, (e) myeloid dendritic cells, (f) T cells, (g) T cells, (h) CD4^+^ T cells and (i) CD8^+^ T cells. Intracellular staining was performed (l) to identify IFN-γ, IL-4, IL-13 or IL-17 producing CD4^+^ cells. (m) Pro-MMP9 was quantified by ELISA at 14 days post-infection. The data are pooled from at least two independent experiments (n=2-7). ns not significant, *p<0.05, **p<0.01, ***p<0.001 compared to control (Mann-Whitney t test).

We also found a selective enrichment of CD4^+^ IL-17A^+^ T cell, but not of CD4^+^ IL-4^+^, CD4^+^ IFN-γ^+^ or CD4+ IL-13^+^ T cells (Figure 3 l). On the contrary, CD8^+^ T cells capable of producing IFN-γ, IL-17A, IL-4 or IL-13 were few and did not change over the course of chronic NTHi infection (data not shown). We quantified the levels of pro-MMP9 as a marker associated to tissue remodeling processes [24]. Levels of pro-MMP-9 were significantly upregulated during chronic NTHi infection in the lung (Figure 3 m), confirming that NTHi persistence is associated with sustained tissue remodeling processes. Thus, all together these data indicate that NTHi persistence in the lower airways induces local Type 17 immune manifestations and sustained tissue remodeling processes.

### Pathogenicity of a CF clinical isolate during persistence in murine airways

We tested whether a CF NTHi isolate may display a worse pathogenic potential in comparison to the reference laboratory strain at the advanced phase of chronic lung colonization. We exploited our new model of NTHi lung persistence in order to compare bacterial burdens and host response between the reference laboratory strain and a CF isolate, named “NTHi50”. As shown (Figure 4 a, b), NTHi50 did not differ from the 49766 reference strain in terms of bacterial burden and incidence of chronic infection in C57BL/6N mice 14 days post-challenge. We then investigated whether the inflammatory response in terms of quality and burden could be different during the persistence of those NTHi strains. We evaluated both histological lesions by the analysis of lung tissue slices stained with H&E (Figure 4 c, d, e) and lung infiltrating cells by FACS analysis in the lung. Both analyses revealed that the persistence of NTHi50 induced a host response similar to that due to the 49766 strain. As a matter of fact, the sustained immune response was characterized by T cells, comprising both γδ^+^ and CD4^+^ subsets, and associated with sustained neutrophil recruitment (Figure 4 f-m). In addition, by immunohistochemistry, we found that CD3^+^ cells were localized in lymphocyte aggregates (Figure 5 and S2) at the advanced stage of chronic infection with both strains. Moreover, we determined that CD3^+^ aggregates were characterized by the strong co-localization of B220 and CXCL13-expressing cells, demonstrating the presence of BALT-like structures at day 14.

**FIGURE 4.**
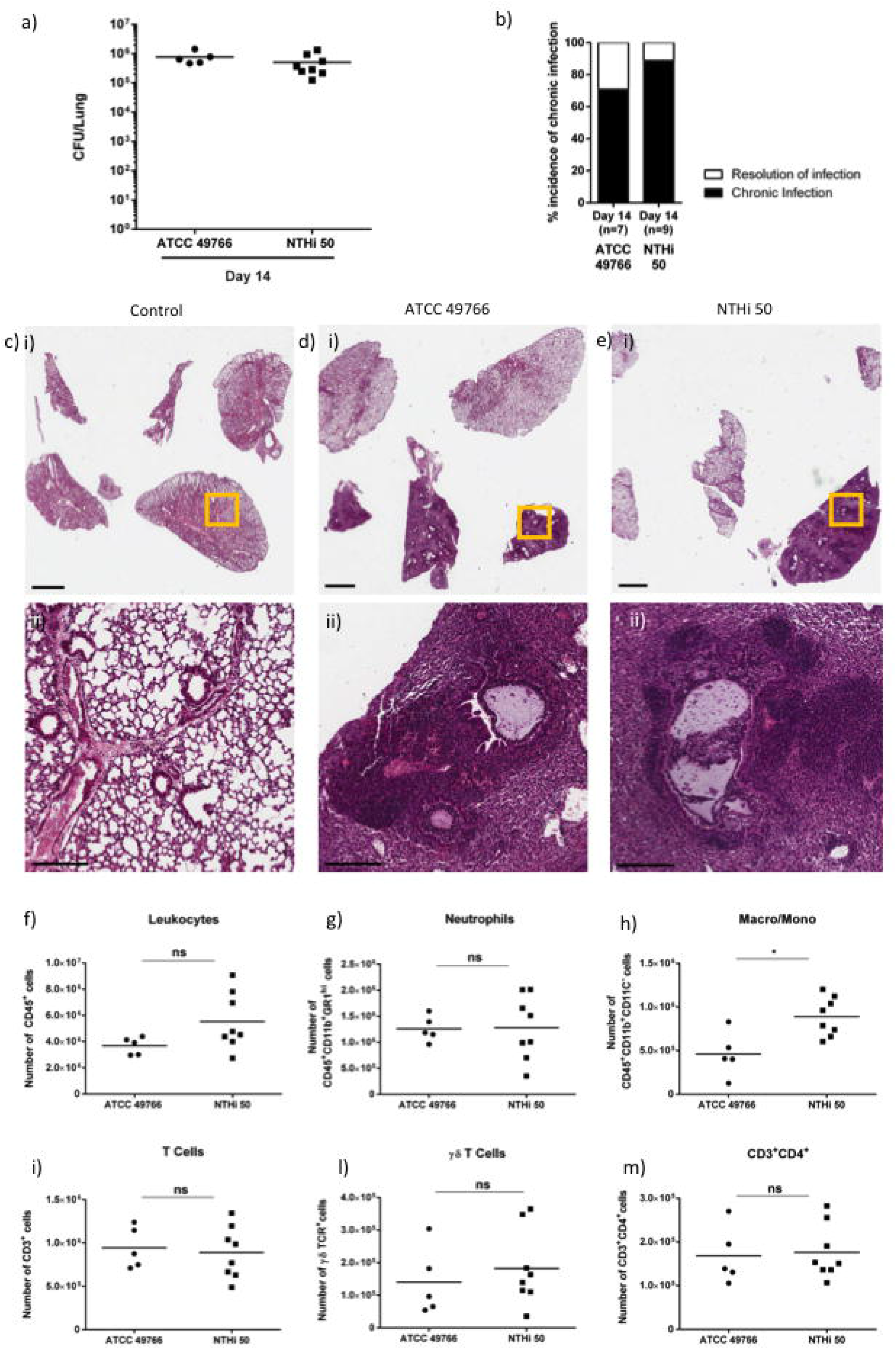
Bacterial burdens, histopathology and immune response in murine airways during the persistence of NTHi CF strain. Mice were infected intratracheally with NTHi embedded agar beads. Bacterial loads in the lung (a) and the incidence of chronic infection (b) were measured at 14 days post infection. H&E staining of lung tissue sections were performed for control mice (c) ATCC 49766 (d) and NTHi 50 (e) infected mice. Representative images are shown from the i) lower (scale bar=2 mm) to the ii) higher magnfication (scale bar=200 μm). Recruitment of (f) leukocytes, (g) neutrophils, (h) monocytes/macrophages, (i) T cells, (l) T cells and (m) CD4 ^+^ T cells was measured by flow cytometric analysis in cell suspensions of murine lungs at 14 days post infection. The data are pooled from at least two independent experiments (n=2-4). ns not significant, *p<0.05 (Mann-Whitney t test).

**FIGURE 5.**
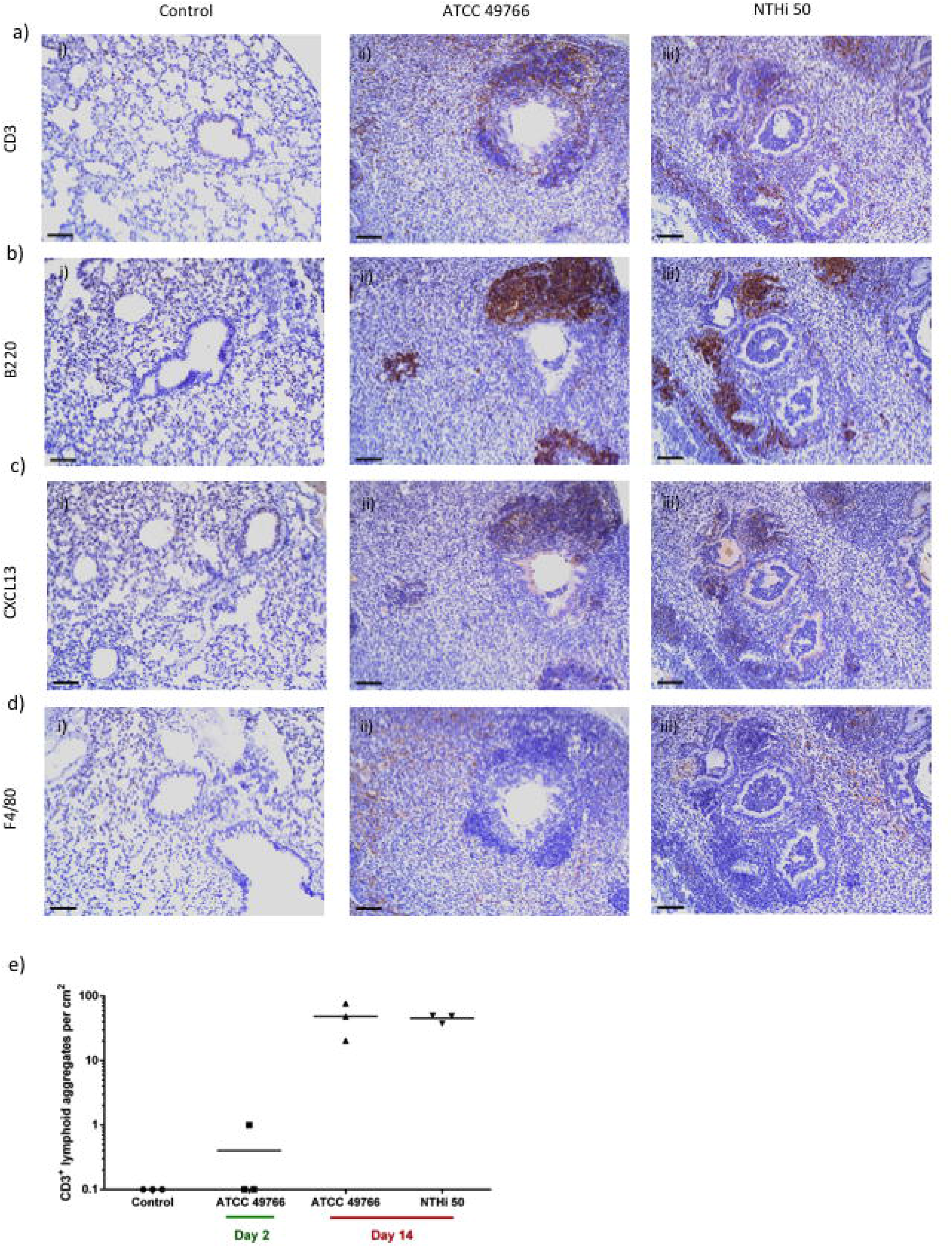
NTHi persistence promotes the formation of peribronchial lymphoid tissue structures. Mice were infected intratracheally with NTHi embedded agar beads and lung immunohistochemistry for the CD3 (a) B220 (b) CXCL13 (c) and F4/80 molecules was performed on lung tissue sections at 14 days post infection for control mice (i), ATCC 49766 (ii) and NTHi 50 (iii) infected mice (scale bar=80 μm). (e) Morphometric analysis of CD3^+^ aggregates in NTHi infected mice. Horizontal bars represent median values. The data are pooled from at least two independent experiments (n=1-2)

To test whether the bacterial pathogenic contribution may differ among CF NTHi isolates, we infected the A549 alveolar epithelial cells with clinical CF isolates (n=8), including NTHi50 and seven other CF clinical strains (Table S1) and evaluated cytotoxicity by analyzing cell viability 6 hours after infection. We observed a similar cytotoxic potential of NTHi clinical strains. As shown in (Figure S1), only NTHi32, NTHi33 and NTHi34 isolates induced a modest (although statistically significant) increase in cell toxicity in comparison to the NTHi reference strain.

Overall, our data highlight that NTHi persistence, rather than NTHi intrinsic virulence, fuels the inflammatory burden in the lungs promoting chronic inflammation characterized by the sustained recruitment of neutrophils and T cells.

## • Discussion

The pathogenic potential of NTHi persistence in CF lung disease is still controversial. Our research question was whether NTHi persistence could be considered as a potential risk factor for lung disease progression by fuelling the inflammatory burden. Our results demonstrate that NTHi colonization is associated with a sustained inflammatory burden, as also indicated by sputum analysis in a small cohort of CF patients. The development of a new murine model was instrumental in mirroring NTHi persistence as observed in clinical settings and, consequently, in overtaking the apparent limited NTHi virulence observed in *in vitro* experiments with bacterial variants. In particular, we found out that NTHi persistence is associated with sustained recruitment of neutrophils in the lung. This pro-inflammatory response seems to be mediated by T cell response (mainly type 17 immunity) and by the presence of BALT in the sub-mucosal lung tissue.

Epidemiological data on *H. influenzae* colonization and disease risk were mainly collected from patients with non-CF bronchiectasis [25–27]. In the context of CF patients with NTHi colonization/infection, only few studies reported on associations with clinical characteristics, morbidity and mortality. *H. influenzae* is more frequently observed in CF patients without a positive history of *P. aeruginosa* lung infection, rather than in those who had been previously colonized at least once [8, 14]. It is worth noting that none of the previous epidemiological studies focused on the contribution of early *H. influenzae* colonization on CF airways inflammation and consequent disease progression. Our data indicate, for the first time, that early colonization by NTHi may account for a higher pro-inflammatory burden.

Interestingly, *H. Influenzae* detection on respiratory tract culture was identified as a risk factor for substantial annual FEV1 decline in 4680 adolescents and young adults with CF from US and Canada within the Epidemiologic Study of CF [28]. However in this large multicentre observational prospective study, it was not possible to ascertain whether this observation was associated with typable *H. influenza* or NTHi subspecies.

To further support our observations in CF patients, prospective observational clinical studies with a higher number of patients are required, possibly in a multicenter study to increase availability of subjects with comparable clinical and demographic characteristics. In this context, the development of biological clinical samples banks (e.g. blood, respiratory samples or clinical strains) associated with a large clinical CF database may be useful to definitively address the potential of early NTHi infection as a risk factor for lung disease progression in CF.

As for clinical and basic science studies, the NTHi literature fails to deeply and fully discriminate persistent infection/colonization from acute/intermittent episodes of infection. The novelty of our work relies not only in the evaluation of CF patients chronically colonized by NTHi, but also in the dissection of the pro-inflammatory potential during NTHi persistence in the airways of mice. To date, animal models have been developed to mimic the acute phase of NTHi infection rather than the chronic phase, limiting steps forward in the field. Airway persistence of NTHi is common in CF disease but also in other lung diseases with different aetiology, including COPD or lung cancer [29–31]. In this context, *Croasdell A. et al*. exploited an acute infection model in which different amounts of NTHi were inoculated by oropharyngeal aspiration in C57BL/6J mice (up to 10^8^ CFU) and 24 hours post-infection they found >90% of bacteria had been cleared, independently from the inoculum dose [32]. To date, mouse models based on the injection of NTHi planktonic bacterial cells in immunocompetent mice are useful for studying the acute infection phase rather than NTHi persistence. In the present study, we were able to evaluate the pro-inflammatory burden caused by NTHi persistence, associated with constitutive recruitment of neutrophils, as well as with increased levels of both γδ^+^ and CD4^+^ T cell subsets in the late phase of bacterial persistence. Previous studies identified a protective role of type 17 immunity (intended as IL-17 secreting cells) during transient and acute NTHi infection in mice [29] [33]. Here, we found that chronic inflammation due to NTHi persistence is associated with CD3^+^ CD4^+^ IL-17^+^ cells. As previously demonstrated for *P. aeruginosa*, we could speculate that type 17 immunity may play a double-edged sword activity during NTHi infection: it could be protective by increasing host resistance in the early phase of infection, while being detrimental during the advanced phase of chronic colonization by increasing the immunopathology [19, 34]. Further studies using transgenic mice or inhibitory drugs could address this hypothesis.

In CF, the pathogenic potential of NTHi persistent infections is unknown, although this bacterium often colonizes CF patients with preserved lung function early in life [8, 10]. Our data suggest that NTHi persistence in mice can induce an inflammatory response similar to those mediated by other well know CF pathogens (e.g. *P. aeruginosa* or *S. aureus*). Here the NTHi-driven peribronchial lymphoid tissue structure associated with sustained neutrophils recruitment, mimics pathological features similar to those observed for longterm *S. aureus* or *P. aeruginosa* colonization in mice or in CF patients with advanced disease [19, 23].

Interestingly, results of long-term infection with laboratory and clinical strains in mice suggest that the pro-inflammatory burden is mainly mediated by the immunopathological responses to NTHi persistence rather than by the specific pathogenic virulence of different bacterial strains. The in vitro results obtained when different clinical isolates were tested further support a similar pathogenicity. Of course, a study considering a higher number of clinical isolates is required to determine whether specific phenotypic traits can affect the pathogenic potential of different NTHi variants.

Overall, these data suggest that NTHi persistence may sustain a pro-inflammatory response similarly to other CF pathogens such as *S. aureus* or *P. aeruginosa*. In this context, the question whether NTHi-inflammatory response may promote the colonization of the airways by other pathogens such as *P. aeruginosa* or *S. aureus* as occurs in CF patients, remains to be addressed. It is desirable that future studies would not exclusively contrast NTHi bacterial growth (e.g. with antimicrobial compounds), but also target to limit the NTHi-induced inflammatory response (e.g. with anti-inflammatory drugs) in CF lung disease. These approaches will help to better explore the contribution of NTHi persistence on disease progression, pulmonary exacerbations or super infection events, such as the “predisposition” to *P. aeruginosa* infection. Moreover, NTHi infections are also commonly associated with other chronic respiratory diseases with different aetiology, such as COPD [3, 4, 27]. Therefore, the murine model developed in this study will be exploitable to determine the contribution of the bacterial persistence and new therapeutic approaches for other lung diseases mediated by NTHi infection.

In conclusion, our results demonstrate that NTHi persistence, common during early lung disease, contributes to the exaggerated lung inflammatory burden and may contribute in declining lung function.

## Acknowledgments

The Authors thank Fiocchi A. (OMC, Mouse Histopathology Unit, San Raffaele Scientific Institute, Milano, Italy) for the mouse histopathology preparations and the San Raffaele Microscopy facility (Alembic) for images acquisition.

This work was supported by OSR_Seed Grant to NIL.

The funders had no role in study design, data collection and analysis, decision to publish, or preparation of the manuscript.

## Author contributions

Conceiving and designing the experiments: N.I.L., C.Ci.; performing experiments: F.S., G.R., D.G. B.S. and N.I.L.; analyzing data: F.S., G.R., B.S., L.C., V.D., C.Ci. and N.I.L.; performing statistical analysis: F.S., G.R., V.D. and N.I.L.; interpretation of the experiments results: F.S., G.R., A.B., L.C., D.C., V.D., C.Co., C.Ci. and N.I.L.; preparing figures: F.S., G.R. and N.I.L.; writing the manuscript: F.S., G.R., D.C., L.C., V.D., A.B. C.Ci. and N.I.L.;

## Notes

### Competing Interest Statement

The authors have declared no competing interest.

