## Supplementary for "The persistence of Nontypeable *Haemophilus influenzae* fuels airway inflammation"

***^3^*** *IRCCS San Raffaele Scientific Institute, Division of Immunology, Transplantation, and Infectious Diseases, Infections and cystic fibrosis unit, Milan, Italy.*

***^4^***  *Cystic Fibrosis Microbiology Laboratory, Fondazione IRCCS Ca' Granda, Milan, Italy.*

***^5^*** *Cystic Fibrosis Regional Reference Center, Fondazione IRCCS Ca’ Granda, Ospedale Maggiore Policlinico, University of Milan Milano, Italy*

Running title: Nontypeable Haemophilus influenzae persistence and inflammation

Keywords: NTHi, inflammation, respiratory infection, il-17, mouse model.

This article has Supplementary Information.

*To whom correspondence should be addressed:

**Supplementary methods**

**Inclusion and exclusion criteria for CF patients**

Inclusion criteria were: a) diagnosis after neonatal screening or after onset of symptoms and followed since birth in the centre; b) complete clinical history available in the computerized database; c) at least two visits yearly since diagnosis; d) availability of at least four yearly sputum cultures for microbiological ascertainment; e) at least one CT every year; f) at least one respiratory function test/year after the age of 4.8 years. **%**FEV_1_ measurements were performed in routine care visits and data were collected accordingly with the local ethics committee.
Exclusion criteria were: i) positive culture for other CF pathogens*;* ii) acute pulmonary exacerbation.

**Agar beads production and NTHi lung infection**

NTHi strains were cultured in brain heart infusion (BHI, BD) supplemented with 10 µg haemin mL^-1^ and 5 µg NAD mL^-1^(sBHI) overnight at 37°C, adjusted to a starting OD^600^ of 0.1, grown to mid-log phase (3h) at 37°C 200 RPM, to allow agar bead preparation. Then, the bacteria were harvested by centrifugation and resuspended in 1 mL of phosphate-buffered saline (PBS; pH 7.4). Bacteria were added to 9 ml of sBHI agar, previously warmed to 45°C. This mixture was pipetted forcefully into 150 ml of heavy mineral oil at 45°C and stirred rapidly with a magnetic stirring bar for 6 min at room temperature, followed by cooling at 4°C for 35 min with a slow and continuous stirring. The oil-agar mixture was centrifuged at 4,000 rpm for 15 min to sediment the beads, which were washed six times in PBS. The number of NTHi colony forming units (CFU) in the beads was determined by plating serial dilutions of the homogenized bacteria-bead suspension on sBHI agar plates. The inoculum was prepared by diluting the beads suspension with PBS to 1 × 10^7^ CFU per inoculum. Immunocompetent C57BL/6NCrlBR male mice (8-10 weeks of age) were purchased from Charles River (Calco, Italy), shipped in protective, filtered containers, transported in climate-controlled trucks, and allowed to acclimatize for at least two days in the animal house prior to use. Mice were maintained in the biosafety level 3 (BSL-3) facility at San Raffaele Scientific Institute (Milano, Italia) where 3-5 mice per cage were housed. Mice were maintained in sterile ventilated cages. Mice were fed with standard rodent autoclaved chow (VRFI, Special Diets Services, UK) and autoclaved tap water. Fluorescent lights were cycled 12h on, 12h off, and ambient temperature (23±1°C) and relative humidity (40-60%) were regulated. For infection experiments, mice were anesthetized by an intraperitoneal injection of a solution of ketamine (100 mg/kg) and xylazinein (10 mg/kg) in 0.9% NaCl and administered at a volume of 0.015 ml/g body weight. Mice were placed in a supine position. The trachea was directly visualised by ventral midline, exposed and intubated with a sterile, flexible 22-g cannula attached to a 1 ml syringe. An inoculum of 50 μl of agar-bead suspension was implanted via the cannula into the lung. After inoculation, all incisions were closed by suture. Mice were monitored daily for coat quality, posture, attitude, ambulation, hydration status and body weight. Mice that lost >20% body weight and had evidence of severe clinical disease, such as scruffy coat, inactivity, loss of appetite, poor locomotion, or painful posture, were sacrificed before the termination of the experiments with an overdose of carbon dioxide. Gross lung pathology was noted. Lungs were excised aseptically and homogenized in 2 ml PBS added with protease inhibitors using the homogenizer gentleMACS^TM^ Octo Dissociator. One-hundred μl of the homogenates and 10-fold serial dilutions were spotted onto sBHI. CFU were determined after overnight growth at 37°C.

Infections, treatments and sacrifices in the chronic infection models were all performed in the late morning. In addition, in all the experiments, mice had been subdivided according to the body weight to have similar mean in all the groups of treatment. Animal studies were conducted according to protocols approved by San Raffaele Scientific Institute (Milan, Italy) Institutional Animal Care and Use Committee (IACUC #920) and adhered strictly to the Italian Ministry of Health guidelines for the use and care of experimental animals.

**NTHi infection in A549 cell line**

Human alveolar epithelial A549 cells were cultured in Dulbecco Modified Eagle Medium (DMEM, Lonza) supplemented with 10% FBS, 2 nM L-glutamine and 100 U/ml penicillin and 100 µg/µl streptomycin. Cells were seeded in flat-bottom 96-well plates at a density of 2x10^4^ cells per well in 100 µL of DMEM. NTHi was grown on brain heart infusion (BHI, BD) agar plates supplemented with 10 µg haemin mL^-1^ and 5 µg NAD mL^-1^ (sBHI) at 37 °C in 5% CO2 overnight, then harvested and incubated overnight in sBHI broth. Next, bacteria were subcultured to log phase growth (3h) and resuspended in PBS to the desired CFUs. Then, 10 µL of bacteria suspension per well were inoculated onto the cells at the MOI of 100. Infected cells were incubated for 6 hours.

**Supplementary Tables**

| **SUPPLEMENTARY TABLE 1 NTHi clinical strains** | |  |  |  |  |
| --- | --- | --- | --- | --- | --- |
| **Strain name** | **Age** | | **Bacterial load** | **Mutation** | **Other colonizations** |
| **NTHi 31** | 5 | | 1,00E+05 | W1282X/R117H (7/7) | - |
| **NTHi 32** | 9 | | 1,00E+05 | DF508M/G542X | STAA/L+ |
| **NTHi 33** | 39 | | 1,00E+06 | DF508/DF508 | STAA |
| **NTHi 34** | 5 | | 1,00E+06 | DF508/R117G | PNU |
| **NTHi 39** | 9 | | 1,00E+05 | DF508/DF508 | - |
| **NTHi 46** | 6 | | 1,00E+05 | DF508/DF508 | - |
| **NTHi 47** | 40 | | 1,00E+06 | DF508/POLI T5 TG12 | STAA |
| **NTHi 50** | 4 | | 1,00E+05 | DF508/W1282X | - |

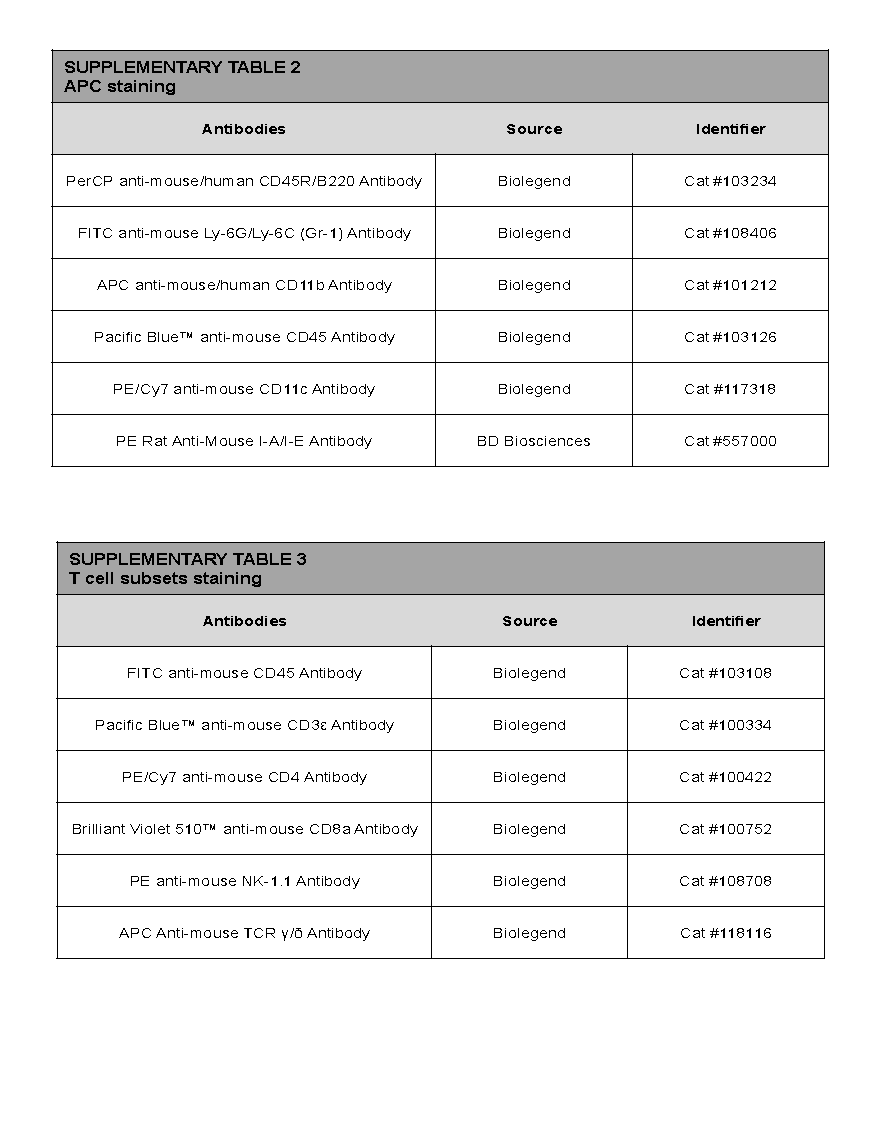

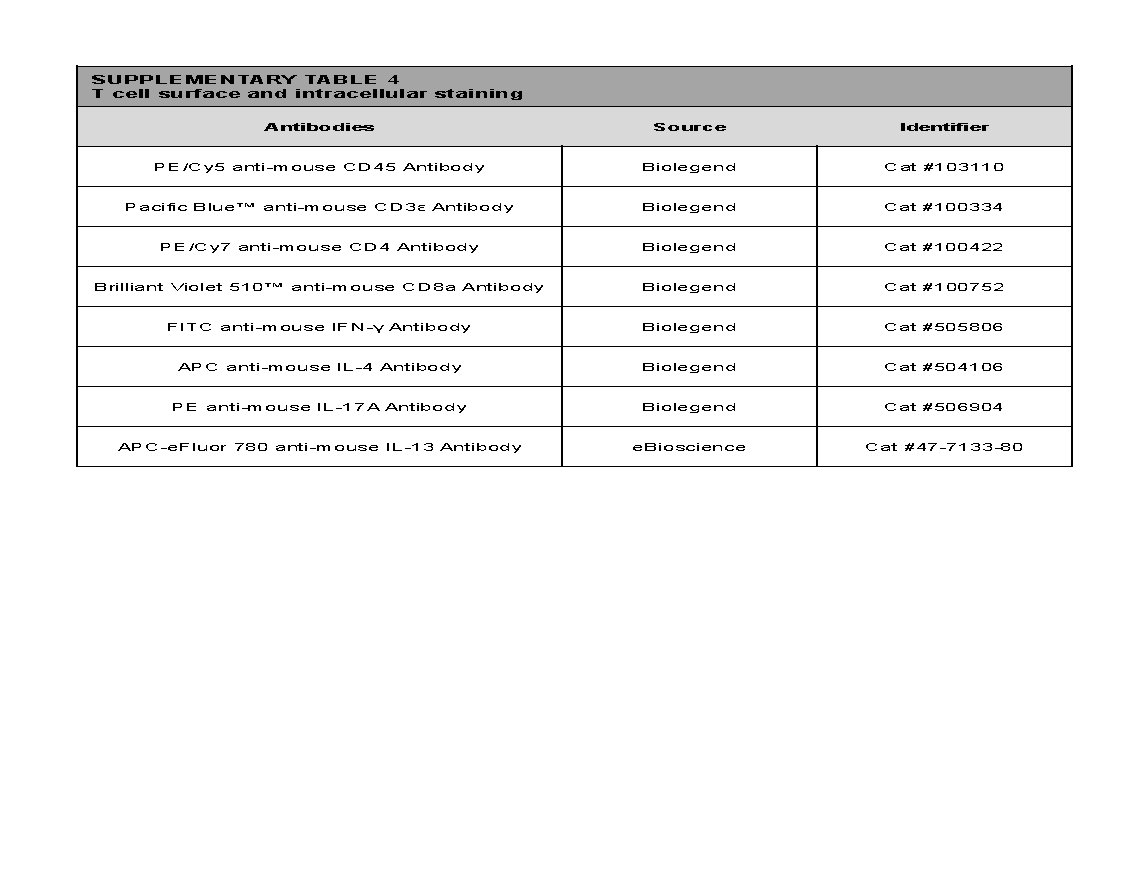

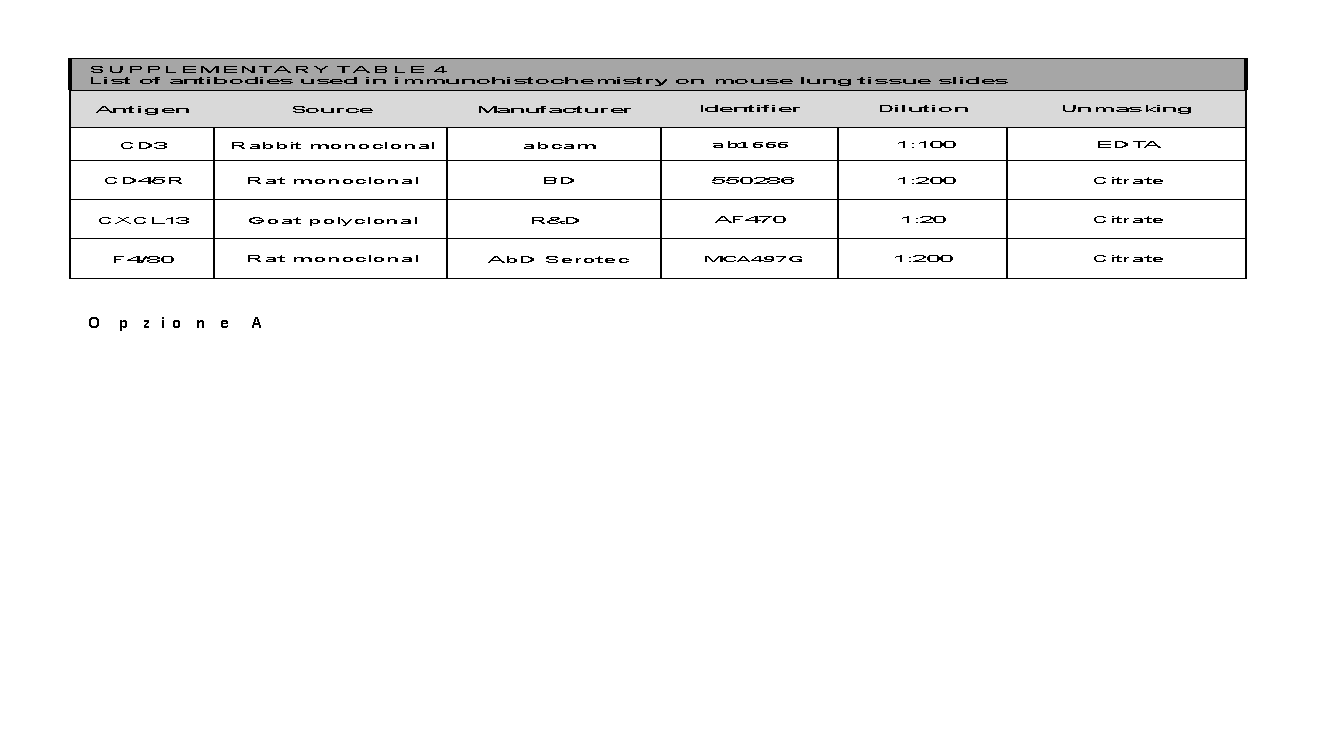

**Supplementary Figures**

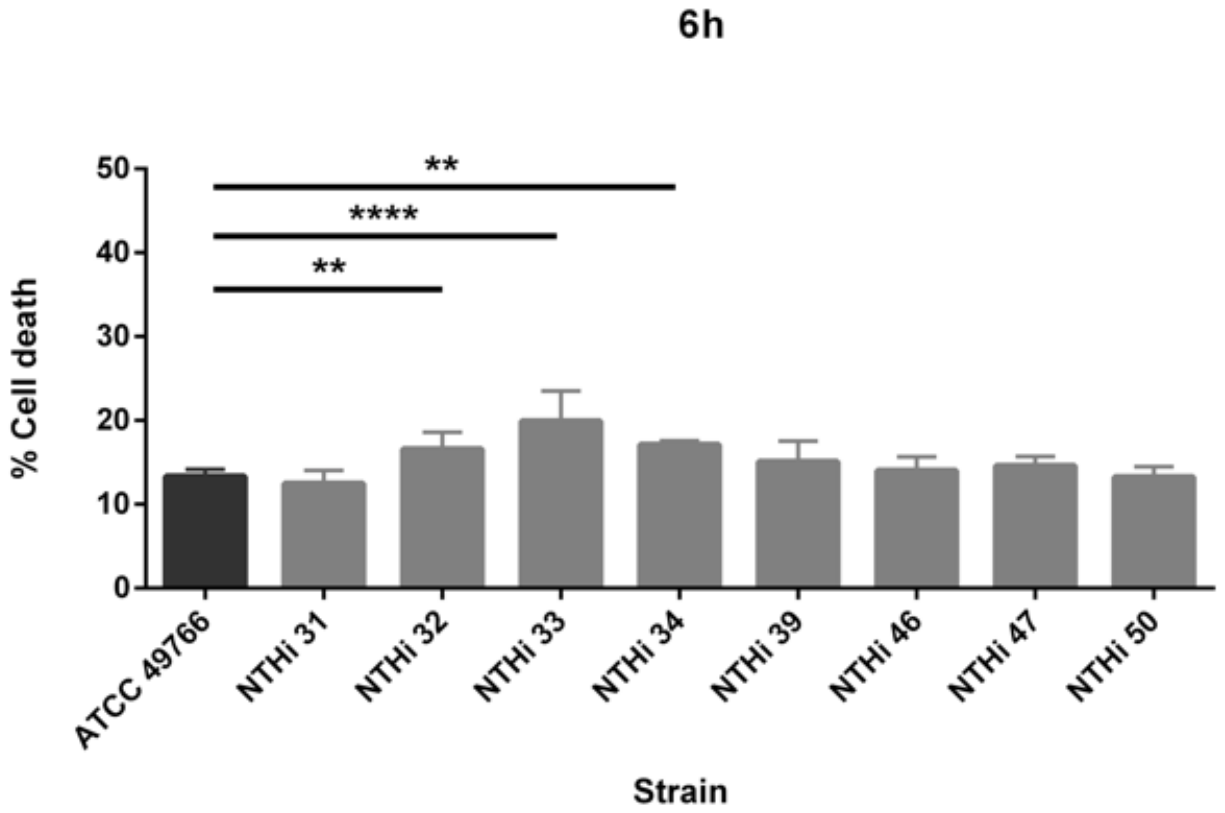

FIGURE S1 A549 cells were infected with eight different clinical strains of NTHi plus the reference strain ATCC49766 at a MOI of 100. Percentage of cell death was assessed at 6 hours post infection by LDH assay. Data are presented as mean ± SD. Ordinary 2-way ANOVA and Bonferroni multiple comparisons test versus ATCC 49766 has been performed as statistical analysis. The data are pooled from at least three independent experiments. ** = p<0.01; **** = p<0.0001.

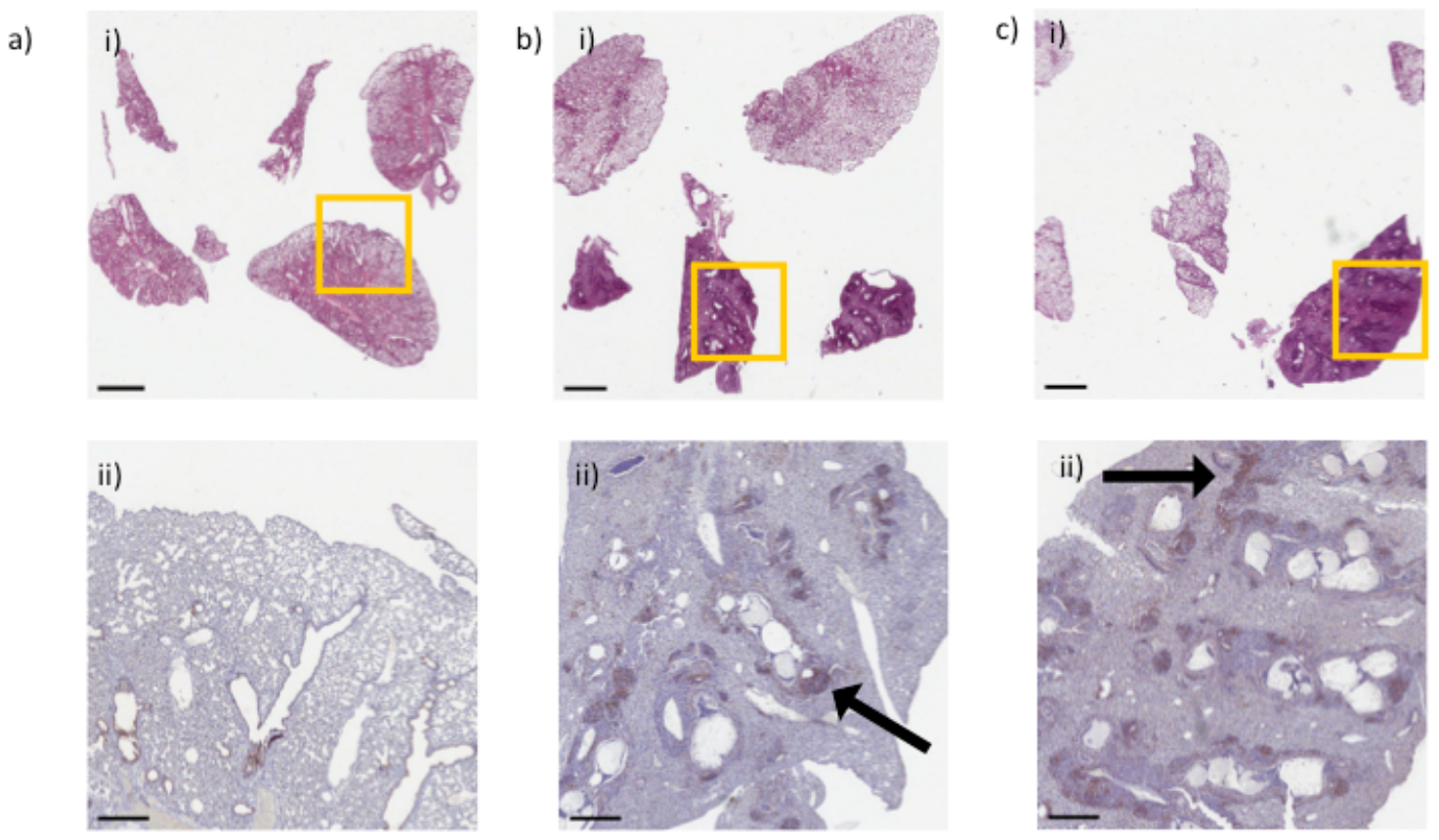

FIGURE S2 NTHi persistence promotes the formation of lymphoid aggregates. Mice were infected intratracheally with NTHi embedded agar beads. H&E stainings (i) and immunohistochemistry for the CD3 molecule (ii) of lung tissue sections were performed at 14 days post infection for control mice (a) ATCC 49766 (b) and NTHi 50 (c) infected mice. Representative images are shown from the lower i) to the higher (H&E, scale bar=2 mm); ii) magnification (CD3+, scale bar=200 µm). Some BALT-like structures are indicated by arrows. The data are pooled from at least two independent experiments (n=1-2)
